# A benchmarking of workflows for detecting differential splicing and differential expression at isoform level in human RNA-seq studies

**DOI:** 10.1101/156752

**Authors:** Gabriela A. Merino, Ana Conesa, Elmer A. Fernández

**Affiliations:** Centro de Investigación y Desarrollo en Inmunología y Enfermedades Infecciosas (CIDIE), CONICET, Av. Armada Argentina 3555, X5016DHK Córdoba, Argentina; Universidad Nacional de Córdoba, Facultad de Ciencias Exactas Físicas y Naturales, Av. Vélez Sarsfield 1611, X5016GCA Córdoba, Argentina; Genomics of Gene Expression Laboratory, Centro de Investigación Príncipe Felipe, Avda. Eduardo Primo Yúfera 3, 46020 Valencia, Spain; Microbiology and Cell Science Department, Institute for Food and Agricultural Research, University of Florida, Gainesville, Florida, USA

**Author notes:** **To whom correspondence should be addressed.** Elmer A. Fernández.

**Keywords:** Alternative Splicing, Differential Expression, RNA-seq

## Abstract

Over the last few years, RNA-seq has been used to study alterations in alternative splicing related to several diseases. Bioinformatics workflows used to perform these studies can be divided into two groups, those finding changes in the absolute isoform expression and those studying differential splicing. Many computational methods for transcriptomics analysis have been developed, evaluated and compared; however, there are not enough reports of systematic and objective assessment of processing pipelines as a whole. Moreover, comparative studies have been performed considering separately the changes in absolute or relative isoform expression levels. Consequently, no consensus exists about the best practices and appropriate workflows to analyse alternative and differential splicing. To assist the adequate pipeline choice, we present here a benchmarking of nine commonly used workflows to detect differential isoform expression and splicing. We evaluated the workflows performance over three different experimental scenarios where changes in absolute and relative isoform expression occurred simultaneously. In addition, the effect of the number of isoforms per gene, and the magnitude of the expression change over pipeline performances were also evaluated. Our results suggest that workflow performance is influenced by the number of replicates per condition and the conditions heterogeneity. In general, workflows based on DESeq, DEXSeq, Limma and NOISeq performed well over a wide range of transcriptomics experiments. In particular, we suggest the use of workflows based on Limma when high precision is required, and DESeq2 and DEXseq pipelines to prioritize sensitivity. When several replicates per condition are available, NOISeq and Limma pipelines are indicated.

## INTRODUCTION

In high eukaryotes, many genes can produce multiple transcripts through alternative splicing (AS), a post-transcriptional regulatory mechanism responsible for the functional complexity and protein diversity made from a small number of genes [1, 2]. Splicing patterns are constantly changing, allowing organisms to respond to modifications in their environment [3, 4]. For instance, more than 90% of human genes are naturally alternatively spliced and misregulations of AS causing changes in absolute or relative isoform expression have been related to several diseases, including cancer [5]. Hence, the determination of changes in splicing patterns is an important issue in basic and applied biomedical research. Today, RNA-seq is the most widely used technique to analyse transcriptome expression dynamics, including AS [6].

In the analysis of AS, two types of changes in isoform expression can be envisioned: *Differential Isoform Expression* (DIE) and *Differential Splicing* (DS). DIE refers to a change in the absolute expression of an isoform, whereas DS is related to changes in isoform proportions [7]. In both cases, the transcriptomic analysis is based on quantification at different levels (i.e. isoform, exon) than gene expression [6]. Several works have been published comparing and evaluating isoform quantification methods using synthetic and/or real RNA-seq data [6-9]. Moreover, numerous differential expression (DE) analysis tools exist for the study of DIE and DS, generally in a separate way [10-13]. In general, specific methods have been developed for DS analysis while DE methods at the gene level have been applied to the study of DIE [10]. Although those methods are well-known in DE analysis at the gene level, their performances over isoform expression data have not been deeply evaluated. Complementary, while some studies comparing methods to detect changes in AS have been published, they are mainly based on a descriptive characterization of method features [1, 7, 14-15]. Hence, a systematic evaluation of workflow performance is needed to further assist the choice of the appropriate set of tools for AS among a plethora of available methods. In this sense, the most complete reported work compares eight popular software tools using both simulated and real RNA-seq data in several scenarios [16]. However, this work focus only on DS changes without consideration of DIE, or DIE and DS occurring simultaneously, and only uses plant data. Thus, there is no clear consensus about the best practices or workflows that should be used or combined to obtain a comprehensive assessment of AS changes in human RNA-seq data involving both DIE and DS together.

Here we present a systematic evaluation and comparison of nine pipelines for the detection of DIE and DS events. In particular, the evaluated DIE workflows were based on isoform expression profiles and used five of the most popular tools: Cuffdiff2 [11], and the R packages: DESeq2 [17], EBSeq [10], Limma [18] and NOISeq [19]. On the other hand, the DS evaluated pipelines were based on Cuffdiff2, and the SplicingCompass [12], DEXSeq [13], and Limma R packages. The study was performed using synthetic RNA-seq datasets where isoform expression profiles were modified and controlled to simulate AS changes based on a real human RNA-seq experiment. The proposed workflows were evaluated in several experimental scenarios, varying the number of genes simulated as differentially expressed, as well as, the number of replicates per conditions. Several performance measures, useful for workflows’ comparison, were obtained. General and practical guidelines based on the number of replicates, sensibility, precision and percentages of true positives are provided in order to aid scientists in the selection of the most appropriate workflows for their data and analysis goals.

## METHODS

### Definition of expression changes at the isoform level

Let us suppose that there are three experimental conditions, A, B, and C, and a gene *g* having two isoforms, *gI* and *gII*, having the expression values listed in Table 1. The comparison of A and B conditions reveals changes in *gI* and *gII* absolute expression,without modifications in their proportions, which is an example of DIE. Note that DIE refers to absolute changes in isoform expression and hence DIE methods use count matrices at the transcript level. When conditions A and C are compared, significant changes in isoform proportions involving small changes in absolute expressions are present. This comparison reveals alterations in the AS mechanism in C respect to A condition, a phenomenon known as DS. The changes in the proportion of the isoforms from the same gene are usually evaluated measuring the changes in the gene’s exon usage.

**Table 1:**
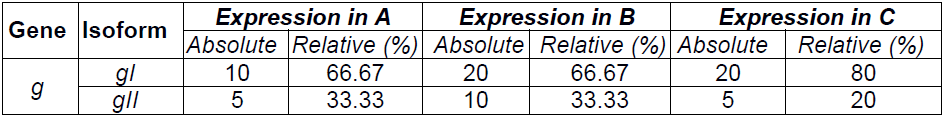
Illustration of changes in absolute and relative isoform expression occurred across three experimental conditions. The comparison of condition A and B reflects the occurrence of differential absolute expression, keeping relative isoform proportions. The comparison of condition B and C reflects alterations in the alternative splicing mechanism causing significant changes in isoform proportions.

### Workflows for differential expression analysis

Seven commonly used methods for DE analysis based on different approaches were chosen to analyse DIE and DS. The selected methods were: EBSeq, DESeq2, NOISeq, SplicingCompass, Limma, DEXSeq and Cuffdiff2. Specific pipelines for them were designed (see Figure 1). The evaluated workflows were called: Cufflinks, DESeq2, EBSeq, Limma and NOISeq, in the case of DIE analysis (solid arrows), and CufflinksDS, DEXSeq, LimmaDS, and SplicingCompass, for DS study (dashed arrows). It is worth nothing, that only Cuffdiff2 and Limma DE tools are able to perform the analysis of both DIE and DS.

**Figure 1:**
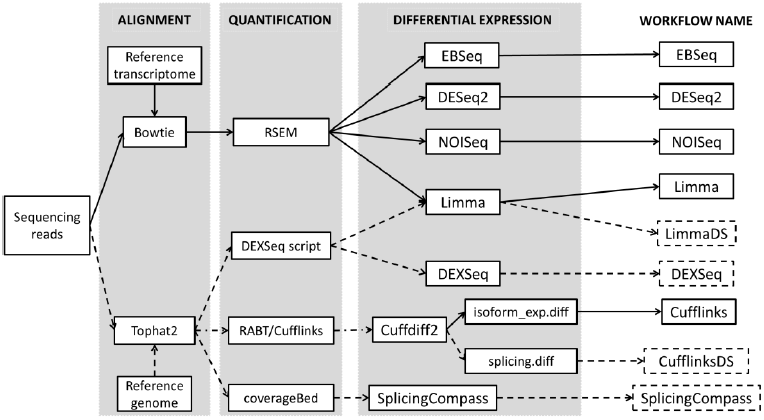
Schema of the nine pipelines evaluated on this work. Five workflows to evaluate differential isoform expression (DIE, solid arrows) and four to analyze differential splicing (DS, dashed arrows) were included. Pipelines were designed following the author’s recommendations to evaluate case-control experiments. All the workflows take as input the sequencing reads and generate a list of isoforms (DIE methods) or genes (DS methods) with significant changes.

### DIE workflows

This group of pipelines takes as input data isoform expression levels obtained by quantification methods based on probabilistic isoform resolution models. These models try to assign reads or fragments to the isoforms they came from modelling the uncertainty derived from multiple isoforms having overlapping sequences [16]. In this work, RSEM [20] was used as a quantification tool to generate isoform count matrices from reads aligned against the human reference transcriptome using the Bowtie tool [21], as suggested by Teng et al and Liu et al [8, 16]. To evaluate DIE, four methods were used i.e. DESeq2, EBSeq, Limma and NOISeq, which are R packages that accept count data at isoform level. EBSeq and DESeq2 assume that the raw expression counts follow a negative binomial distribution, whereas Limma assumes that the logarithmic transformation of expression counts follows a normal distribution. To infer DE changes between experimental conditions, EBSeq uses a Bayesian hierarchical model [10], while DESeq2 combines empirical Bayes shrinkage with generalized linear model estimations to obtain model coefficients and then uses the Wald statistic [17]. The `voom` transformation [22] applies a generalized least squares approach by modelling the mean-variance relationship with precision weights, allowing the use of the classical `eBayes` Limma method to detect the isoform expression changes [18]. The three methods return an FDR adjusted p-value, used here to call DIEs. The `NOISeqbio` tool, from the NOISeq package, is a non-parametric and data adaptive method that uses fold changes and absolute expression differences between the experimental conditions to obtain one statistic per isoform. This method performs a permutation step to obtain the noise distribution, against which the isoform statistics will be compared [19]. `NOISeqbio` returns for each isoform the probability of being differentially expressed (p_de_) and the adjusted p-value is 1-p_de_.

### DS workflows

In the case of DS workflows, the analysis is performed over expression matrices at several levels obtained from alignments against the reference genome. The gapped aligner TopHat2 [23] was used to do this mapping and evaluated DS methods were three R packages, SplicingCompass, DEXSeq, and Limma, and the program Cuffdiff2. The `coverageBed` [24] was used to obtain expression matrices for SplicingCompass, applying a union transcript model for each gene. With this information, SplicingCompass constructs vectors of exon and junction counts for each gene and sample, then calculates pairwise geometric angles between two samples and uses a t-test to compare geometric angles [12]. DEXSeq is based on negative binomial generalized linear models [13], like DESeq2. Count matrices at exon level for DEXSeq and Limma packages were obtained using the python script provided by the DEXSeq package, disabling the aggregate options, as suggested by Soneson et.al. [7]. DEXSeq and Limma models incorporate an interaction term between the condition and the exon identifier to evaluate changes in the proportion of that exon within a gene and between conditions. The `initSigGenesFromResults` (SplicingCompas), `perGeneQValue` (DEXSeq) and `diffSplice` (Limma) functions were used to compute per gene adjusted p-values.

The C++ Cufflinks2 program was used to calculate the isoform expression values as Fragments Per Kilobase Million (FPKM) from reads aligned to genome sequences [11, 25]. Then, Cuffdiff2 performed DE analysis at isoform and splicing levels, generating the output files of the two workflows: Cufflinks (DIE case) and CufflinksDS (DS case).

In all workflows, significant isoform/gene changes were identified using an adjusted p-value threshold of 0.05. The program versions, as well as all the scripts used in this study, are available in supplementary material.

### Simulated RNA-seq datasets

A replicated human prostate cancer RNA-seq dataset (GSE22260) was used as the reference to generate synthetic data. This dataset consists of 30 samples, ten from normal tissue (control, condition-C) and 20 from prostate carcinoma (tumor, condition-T), sequenced using the Illumina GAII platform with a pair-end protocol. The ten T-C pairs matched samples were discarded to avoid subject correlation. In addition, four samples were tagged as outlier samples by our quality control pipeline [26] and discarded to finally keep 16 samples, eight per condition, that were then used to feed the simulator. Three possible experimental scenarios (S1, S2, and S3) combining DIE and DS events were designed. The S1 and S2 scenarios involved eight non-matched samples, four per each condition. In S1, 5% of total genes were simulated to have expression differences (DIE/DS); whereas, S2 had 10% of changing genes incrementing the between-conditions heterogeneity. The S3 scenario considered the effect of a different number of replicates per condition, involving 16 samples, eight for each condition, with 10% of differentially expressed genes, the same as S2.

A simulation procedure was designed to obtain raw sequencing reads for each subject with controlled DIE and DS. The simulation was based on the `rsem-simulate-reads` tool from RSEM, which takes a customized isoform expression profile and model parameters, computed from a real RNA-seq dataset, to generate synthetic sequencing reads. The implemented simulation procedure is shown in Figure 2 and consists of three steps. In step one, each real sample was aligned against the reference transcriptome and, isoform expression profiles together with RNA-seq model parameters were obtained. In the second step, the expression matrix was pre-processed to exclude low expressed genes (zero counts in at least one replicate of C and T). Then, a set of well-expressed genes (20 counts per million in at least one replicate of C) were randomly chosen to simulate DIE/DS. The expression counts for the *i-th* isoform of the *g-th* gene from the *k-th* condition were modelled by a negative binomial (NB) distribution, *y*_*igk*_ ∼*NB(μ*_*igk*_, *θ*_*igk*_). The sample mean (*μ*_*igk*_) and shape (*θ*_*igk*_) for the NB distribution were computed over condition C and taken as a reference to compute the simulated parameters for both C and T conditions, incorporating DIE/DS in the group of genes to be simulated as differentially expressed. In the third step, the isoform counts for each replicate were generated from NB distributions with the modified mean and shape parameters, obtaining the simulation expression profiles. Finally, transcripts per million for each sample were computed and used to call the `rsem-simulate-reads` function to generate simulated raw reads. In order to provide statistical power to evaluate workflows performance, the third step of the simulation pipeline was run ten times to obtain replications of each scenario keeping the same differentially expressed genes and NB parameters.

**Figure 2:**
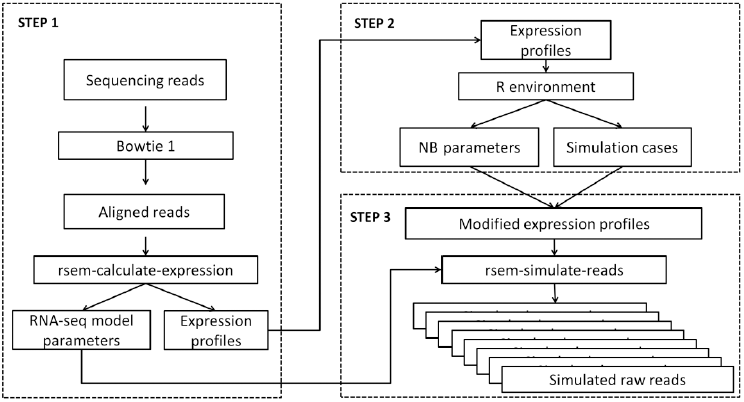
Simulation procedure designed to generate the RNA-seq datasets used to workflows evaluation. The procedure was divided into three steps. The first step was run once for each sample and it started by aligning reads to the reference. Next, alignments were processed to obtain the real isoform expression profiles and RNA-seq model parameters, for each replicate. In the second step, the mean and shape negative binomial parameters for each isoform in each experimental condition were calculated and modified to simulate expression changes. This step was run once for each scenario. The third step was run ten times per each scenario and consisted in generating the simulated isoform expression profiles using a negative binomial distribution with the modified parameters. Finally, the customized expression matrix and the RNA-seq model, estimated before, were used by RSEM to obtain the simulated sequencing reads for each sample.

Since DIE and DS occur simultaneously, both cases were jointly simulated. The set of genes selected to be differentially expressed was divided into four subsets: DE, DIE, DS, and DIEDS. For the DIE and DE groups, changes in the expression of all isoforms of the gene were simulated, without modification of isoform proportions. The DIE group included genes having more than one annotated isoform; whereas, the DE group involved genes having only one annotated transcript. For the DS group, changes in isoform proportion were simulated, without modifications of the overall gene expression. For each gene, the proportion of the most expressed isoform (major isoform) was controlled and the remaining proportions were equally distributed along their other expressed isoforms. Finally, the DIEDS group, the simultaneous occurrence of DIE and DS was simulated. Even though the DS occurrence could derive in DIE, we would include this group where we control DIE and DS presence. More detailed information about simulation groups, subgroups and the computation of simulated profiles can be found in the supplementary material.

### Performance evaluation

Commonly used performance measures were computed to evaluate workflows results at ten simulations from each scenario [27]. The result of each workflow was either a list of significant differentially expressed isoforms (DI), or genes detected as alternative spliced(ASG). Each isoform or gene detected as differentially expressed was called positive (P), and was classified as true positive (TP) or false positive (FP); whereas, an isoform or gene detected as differentially expressed (negative, N) was classified as true negative (TN) or false negative (FN). In particular, for DIE workflows, those isoforms simulated either as DIE, DE, DIEDS or DS were considered as TPs, since DS could cause DIE; but, only the isoforms simulated as DIE, DE or DIEDS were considered as FNs because we did not control if the simulated DS changes caused DIE. Then, accuracy, sensitivity, precision and the F-score (harmonic mean between sensitivity and precision) were computed. The ability of a workflow to deal with false positives was characterized measuring the FP rate (FPR). We also evaluated the effect of simulation subgroup, i.e. DIEDS-2-0.8-0.3, and the effect of the number of isoforms per gene. In the first case, isoforms and genes were clustered according to their simulation subgroup and the TP rate (TPR) was computed in order to determine if the magnitude of deregulation influenced the DI/ASG detection. In the other case, genes were grouped according to their number of annotated isoforms, i.e. 1, 2-4, 5-9 and more than 9 (>9) transcripts and TPRs per each of those groups were computed. Those numbers correspond to the 33, 66 and 99 percentiles, respectively, of the distribution of the number of isoforms per gene in humans.

## RESULTS AND DISCUSSION

Nine workflows for DIE and DS analysis were compared in this study based on synthetic data, where the true status of each isoform or gene was controlled. Three experimental scenarios, S1, S2, and S3 were designed to evaluate the effect of the percentage of differential genes (S1 and S2) and the number of replicates per condition (S2 and S3).

### Concordance of Differential Expression Results

The concordance of differential expression results was evaluated looking at the number and percentage of detected DI/ASG (P and TP) in the ten replicates run for each scenario. Results for DIE and DS workflows are shown in Figure 3 and summarized in Supplementary Tables **S4** and **S5**. In the case of DIE pipelines, the EBSeq workflow detected the highest amount of DI (>8500) in the three tested scenarios, whereas the Cufflinks pipeline found the lowest values, three times lower than the EBSeq results. However, EBSeq had the lowest number of P found simultaneously in the ten simulations (Figure 3**A**), indicating the poor concordance of this method in all three scenarios. On the other hand, DESeq2 and Limma showed a higher concordance of P (>17%, Figure 3**A**) and TP (> 30% Figure 3**B**) along simulations, especially for S2 and S3, showing that they were more robust than EBSeq. Comparing S1 and S2 scenarios, EBSeq and Cufflinks methods did not show differences in the percentage of TPs. On the contrary, DESeq2, Limma and NOISeq increased this percentage in S2 by approximately 5%. TP percentages were increased by 10% for EBSeq and DESeq2 and only 1% for Cufflinks and Limma from S2 to S3. Meanwhile, NOISeq was the only method that showed the highest percentage of TP detections in S2. The FP percentage (Figure 3**C**) was <5% for all the scenarios and pipelines, indicating the effectiveness of the simulation procedure.

**Figure 3:**
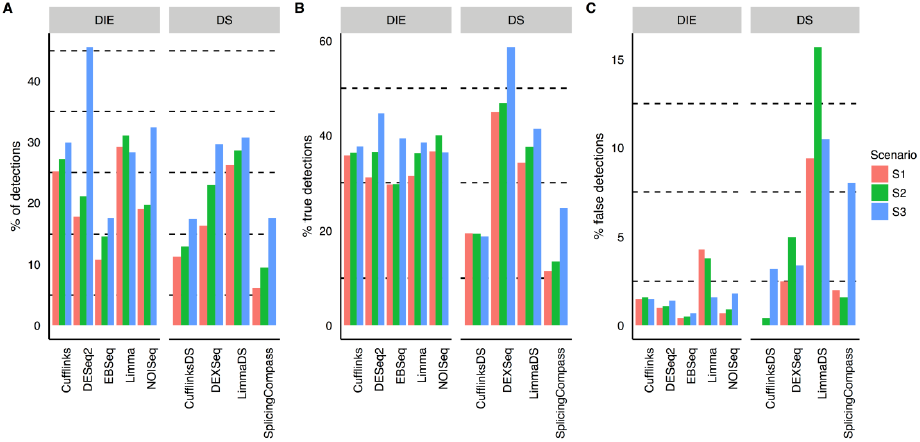
Concordance of workflows results in the ten simulations performed in each scenario. Here, concordance was measured by means of the percentage of significant detections (**A**), true significant detections (**B**) and false significant detections (**C**) of DI/ASG found in the ten runs of each scenario. Each panel is divided into two facets, one for DIE workflows, detecting DI, and other for DS workflows, detecting ASG.

In the case of the DS workflows, CufflinksDS found the lowest average number of ASG (<303); whereas, the highest values were observed for DEXSeq (>423). The lowest and the highest percentage of P were found for SplicingCompass (<20%) and LimmaDS (>25%), respectively (Figure 3**A**). Moreover, SplicingCompass and CufflinksDS had a poor concordance of TP detection (Figure 3**B**). Interestingly all workflows, except CufflinksDS, increased the percentage of P and TP in S2 with respect to S1 and S3. In terms of FP, LimmaDS pipeline showed the highest values, near to 10%.

### Overall performance results

The overall performance measures for the evaluated workflows on the simulated scenarios are listed in Supplementary Table **S6.** All DIE and DS workflows achieved a high accuracy (>0.85) in all scenarios and hence, this measure was not further considered in our comparisons. Sensitivity, precision and F-score for DIE workflows are shown in Figure 4, panels **A-C**. In terms of sensitivity (Figure 4**A**), all DIE pipelines reached values lower than 0.65; the highest values were exhibited for EBSeq (S1 and S3), NOISeq (S2) and DESeq2 (S3), and the lowest for Cufflinks. In terms of precision (Figure 4**B**), EBSeq and NOISeq (S1 and S2) had low performance; whereas, values higher than 0.9 were achieved by Limma, Cufflinks (S2 and S3) and NOISeq (S3). It is worth mentioning that only Limma(S1) and NOISeq (S3) were able to control the imposed FDR, achieving precisions higher than 0.95. When the three experimental scenarios were compared, an improvement from S1 to S3 was noted, except for NOISeq’s sensitivity and Limma’s precision, which showed opposite behaviours. In terms of the F-score, the best values (>0.7) were found for DESeq2, EBSeq, Limma (S3) and NOISeq (S1 and S2). Thus, DESeq2 and EBSeq workflows seem to be adequate to DIE analysis. However, if precision is preferred, Limma and NOISeq are recommended.

**Figure 4:**
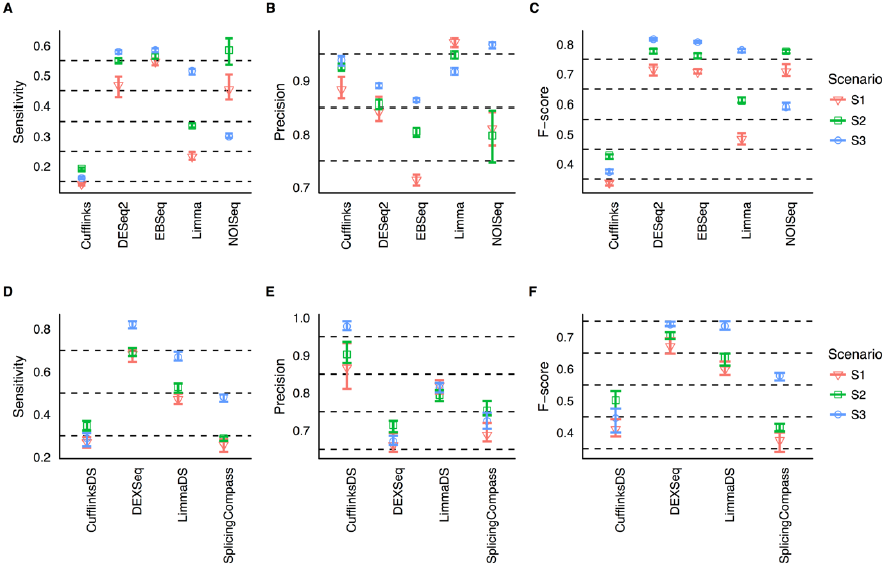
Overall performance measures along ten simulations performed in three experimental scenarios**. A** to **C:** DIE pipelines, **D** to **F:** DS workflows.

The Figure 4**D-F** summarizes the performance results for DS workflows. In terms of sensitivity (Figure 4**D**), DEXSeq and LimmaDS had the best performance, achieving values higher than 0.5 in nearly all scenarios; whereas CufflinksDS and SplicingCompass exhibited the poorest results. However, CufflinksDS showed the highest precision (>0.8), controlling also the FDR in S3 (Figure 4**E**). Although DEXSeq had the lowest precision (<0.75), this method together with LimmaDS, achieved the higher F-score values in all cases (>0.55) and hence these pipelines are adequate to detect ASG with high sensitivity and precision. Particularly, LimmaDS had lower sensitivity than DEXSeq but, it reported ASG more precisely.

The ability to deal with FP results was evaluated using the FPR (Supplementary Figure **S1**). In the case of DIE workflows (Supplementary Figure S1**A**-**C**), the lowest FPR was achieved by Cufflinks in all scenarios, whereas EBSeq, in the three scenarios, and NOISeq, in S1 and S2 had the highest FPR. Regarding DS pipelines (Supplementary Figure S1**D**-**F**), CufflinksDS showed the lowest FP values and DEXSeq the worst. In general, FPR values did not exceed 0.05, with higher values for S2 in comparison with S1. In the S3 all DIE pipelines, except Limma, had lower FPRs in respect to S2.

Based on the poor performances described above for Cufflinks, EBSeq, CufflinksDS and SplicingCompass, these methods were excluded from further analysis and only five pipelines were selected for subsequent evaluations: DESeq2, Limma and NOISeq, for DIE analysis, DEXSeq and LimmaDS for DS study.

### Effect of the number of isoforms

Figure 5 illustrates the relationship between the TPRs and the number of isoforms per gene. Upper (lower) panels show the results for the DIE (DS) workflows and the three evaluated scenarios. For DIE workflows (Figure 5**A-C**), the percentage of TPs was higher for isoforms belonging to genes with only one annotated transcript (gene group “1”) and lower for those belonging to genes with more than nine isoforms (gene group “>9”), in all scenarios. For instance, in S2 (Figure 5**B**) all workflows achieved percentages higher than75% for isoforms from gene group “1”; while the percentage of TPs in gene group “>9” was lower than 50%. We suspected this behavior was caused by lower expression values and complexity of isoform reconstruction process when the number of isoforms per gene increases. DESeq2 and NOISeq showed the highest and similar TPRs in S1 (Figure 5**A**) and S2, while DESeq2 and Limma performed best in S3 (Figure 5**C**). In general, all workflows performed better in S2 compared to S1 and in S3 compared to S2, except for NOISeq, that had poorer TPRs in S3.

**Figure 5:**
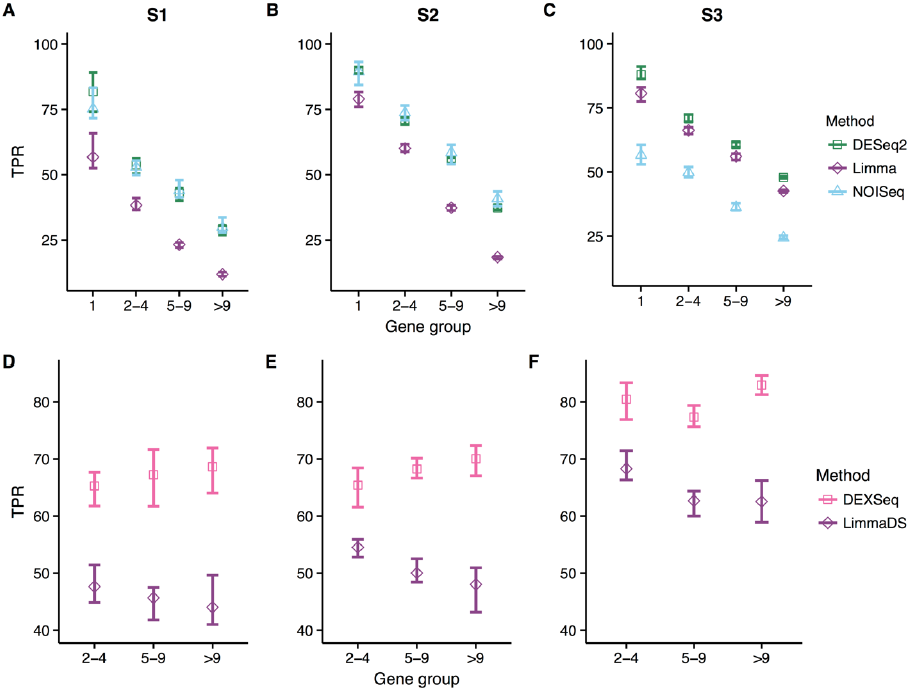
True positive rate of DIE and DS workflows as a function of the number of isoforms per gene. Panels **A**, **B**, **C** are for DIE workflows and **D**, **E**, **F** for DS workflows at S1, S2 and S3, respectively.

In the case of DS workflows (Figure 5, panels **D**, **E**, and **F**) the observed TPRs were very similar in all scenarios and in all gene groups. The highest values were achieved by DEXSeq (> 60%). Notably, LimmaDS showed TPRs higher than 40% that was better than Limma performance in DIE analysis. The TPRs practically did not change between S1 and S2, whereas in S3, DEXSeq and LimmaDS increased the TPRs in all gene groups.

The relationship between FP and the gene groups is illustrated in Supplementary Figure S2. In the case of DIE pipelines, FPs distribution along gene groups was different between scenarios and pipelines. DESeq2 and NOISeq showed similar behavior along S1 and S2, with most of FP (>35%) for gene group “>9”. In addition, the number of FPs increased with the number of isoforms per gene, as expected. Meanwhile, most FPs for Limma were found in the gene group “2-4”. Nevertheless, Limma behaved similarly to DESeq2 and NOISeq in S2 and S3, respectively. For DS workflows, FPs distributions along gene groups and scenarios were similar. In general, FPs were near to 20%, 25% and 45% for gene groups “2-4”, “5-9” and “>9”, respectively. It was observed that for both, DEXSeq and LimmaDS, FPs were more abundant when the number of isoforms per gene increased.

### Effect of the magnitude of differential expression

Finally, the effect of the magnitude of differential expression in the ability of each workflow to detect changes was evaluated (Figure 6). As expected, all DIE pipelines (Figure 6, panels **A**, **B**, and **C**) showed a higher TPR when the magnitude of the expression change was increased, however, differences were evident as a function of the simulation group (DE, DIE, or DIEDS) and the simulation scenario. While all methods had high TPRs when single gene isoforms were simulated with a fold-change of 4 in all scenarios, important differences were observed at fold-changes of 2, where Limma behaved poorly in scenariosS1 and S2 and NOISeq in scenario S3. Surprisingly, all methods had much lower TPRs in the DIE group compared to DE. For example, nearly perfect TP detection was achieved with a 4 fold-change in DE cases, this value dropped to around 50% when talking about DIE transcripts at the same fold-change. This could be explained by the fact that multi-transcripts genes generated both high and low expressed isoforms and all of those were analysed to compute DIE analysis and TPRs calculation. And, as it is known, low expressed transcripts that are differentially expressed are more difficult to detect than those highly expressed. When a change in isoform proportions was included in the simulation group (DIEDS), TPRs were again affected. In scenario S1 all pipelines performed better at detecting true isoform changes when there was also an effect on the relative proportion of the isoform, while this was only the case for transcripts with a 4 fold-change in scenario S3 and was pipeline-dependent in scenario S2. In general, and in agreement with other analyses, Limma performance was comparatively worse at S1 and S2 and NOISeq at S3.

**Figure 6:**
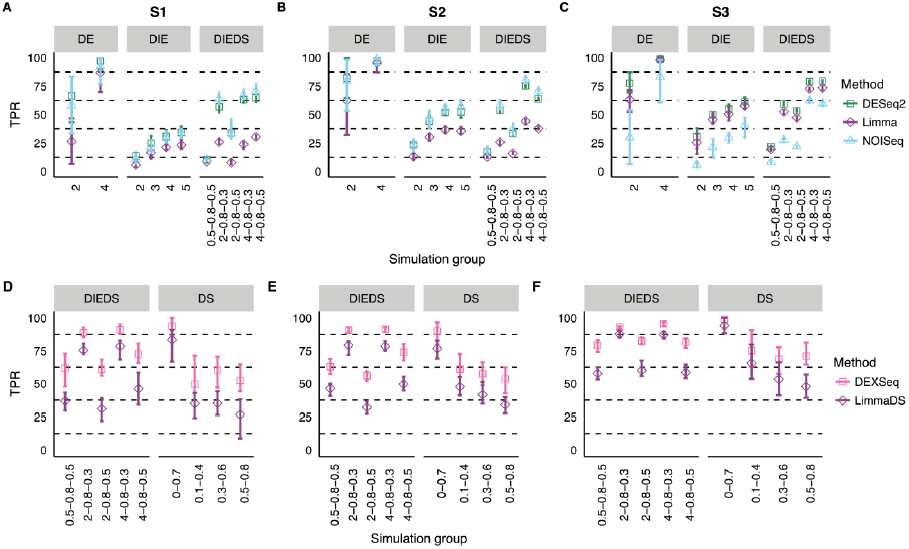
True positive rates (TPR) for DIE and DS workflows as a function of the simulation subgroup as described in Table 2. Panels **A**, **B**, **C** for DIE and **D**, **E**, **F**, for DS workflows at scenarios S1, S2 and S3 respectively.

**Figure 7:**
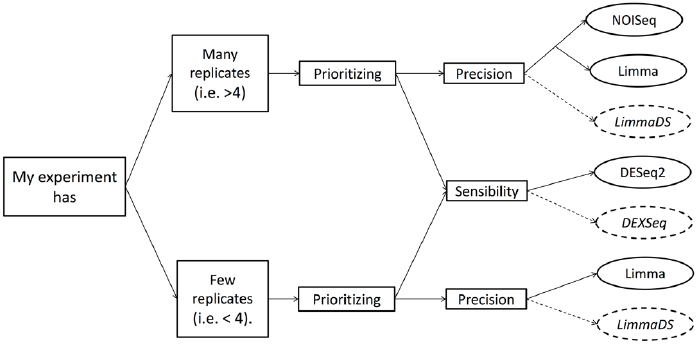
Schema of workflows selection based on the research experiment. Each circle box contains the DIE (solid) and the DS (dashed) pipelines recommended for transcriptomic analysis.

For DS pipelines (Figure 6, panels **D**, **E**, and **F**), results were more predictable. As a general rule, DEXSeq performed better than LimmaDS in all simulation groups and scenarios. In addition, higher TPRs were found when the magnitude of the DS was bigger. In the DIEDS group, good performance was basically associated at the magnitude of the splicing change (values higher than 0.75 for a 0.8-0.3 difference) and to a much lesser extent to the magnitude of the total fold change of the gene (similar results for 0.5, 2 and 4 global gene fold-changes and only slightly lower in the DS 0.5-0.8 subgroup that is zero global gene change). For genes with splicing and not total expression differences (DS), better TPRs were found when the major isoform had zero (DS-0-0.7) or low (DS-0.1-0.4) relative expressionincondition,although the biggest effect for an improved TPR was given by the magnitude of the differential splicing: the DS-0-0.7 subgroup had much higher TPRs than DS-0.1-0.4, DS-0.3-0.6 and DS-0.5-0.8. Both for DIEDS and DS groups overall performances were better when more replicates were present (S3 vs S1 and S2).

**Table 2:** Simulation groups and subgroups. Groups were defined according to the combination of changes in absolute and relative expression of gene isoforms at two experimental conditions.

| *Group* | *Fold Change at isoform expression* | *Change in the proportion of the major isoform* | *Simulation subgroup* |
| --- | --- | --- | --- |
| *DE* | 2 | No change | DE-2 |
|  | 4 | No change | DE-4 |
| *DIE* | 2 | No change | DIE-2 |
|  | 3 | No change | DIE-3 |
|  | 4 | No change | DIE-4 |
|  | 5 | No change | DIE-5 |
| *DS* | No change | 0 to 0.7 | DS-0-0.7 |
|  | No change | 0.1 to 0.4 | DS-0.1-0.4 |
|  | No change | 0.3 to 0.6 | DS-0.3-0.6 |
|  | No change | 0.5 to 0.8 | DS-0.5-0.8 |
| *DIEDS* | 0.5 | 0.8 to 0.5 | DIEDS-0.5-0.8-0.5 |
|  | 2 | 0.8 to 0.3 | DIEDS-2-0.8-0.3 |
|  | 2 | 0.8 to 0.5 | DIEDS-2-0.8-0.5 |
|  | 4 | 0.8 to 0.3 | DIEDS-4-0.8-0.3 |
|  | 4 | 0.8 to 0.5 | DIEDS-4-0.8-0.5 |

DE, Differential Expression; DIE, Differential Isoform Expression; DS, Differential Splicing; DIEDS, Differential Isoform Expression and Differential Splicing

## CONCLUSIONS

In this study, we performed a systematic evaluation of workflows for DIE and DS analysis using simulated RNA-seq datasets based on a real human experiment. The goal of our work was to provide guidelines for choosing appropriate analysis strategies for researchers interested in different modalities of isoform expression changes. For this, we evaluated nine workflows in a variety of expression setups and experimental configurations. We tailored our analysis to human transcriptomics datasets with variability similar to tumor-healthy subject samples. The validity of our study for other types of data (i.e. cell-lines or organisms with lower complexity) remains to be demonstrated.

In general terms, we found that a better scenario for case-control comparisons was when more differential genes (10% vs 5%) and replicates per condition (8 vs 4) were available (S3). For this configuration, we found the highest number of DI/ASG, TPs and concordance among replicated simulation. Best performing workflows were DESeq2, Limma and NOISeq for DIE analysis and DEXSeq and LimmaDS for DS testing.

We used precision, sensitivity and F-score as performance measures. For experiments with a low number of replicates, the best pipelines to DIE analysis were DESeq2 and Limma. Based on our results, we concluded that, if high sensitivity is preferred, DESeq2 is the most indicated, while the Limma pipeline should be used if higher precision is important. For experiments with a large number of replicates, NOISeq is more restrictive and precise than Limma. For DS pipelines, we found that DEXSeq was the best in terms of sensitivity and F-score. However, precision of this method was lower than the one achieved by LimmaDS, which reached the DEXSeq F-score values when the number of replicates was increased. We concluded that these two workflows are indicated for DS analysis, the first one prioritizing sensitivity and the second precision. When the FPR was evaluated, we found that both Limma and LimmaDS workflows were superior to DESeq2 and DEXSeq, respectively.

We also evaluated the effect of the number of isoforms per gene in the percentage of true and false positives. DIE pipelines were found to be more influenced by the number of isoforms per gene than the DS workflows, probably by the presence of low-expressed isoforms. In addition, the TPRs for DIE workflows decreased as the number of isoforms per gene increased. In particular, we found TPR between 30% to 90% using four replicates for DESeq2 and Limma. Whereas, NOISeq reached values between 25% and 60% when more replicates were available. Using DEXSeq or LimmaDS, we found near to 40% of TPs with fewer replicates. Meanwhile, TPR increased to 60% when more replicates were available. We also found that, in DIE and DS cases, most of the false positives were related to genes with more than nine isoforms.

Exploring the effect of the magnitude of differential expression on TPRs, we noted that this was higher for isoforms with greater expression changes, with or without changes in the AS. In the case of DESeq2 and NOISeq, TPs detection was further improved when the percentage of differentially expressed genes was higher. This suggests that these pipelines benefit from an extended regulation in the data and might have problems in detecting differential expression when this affects only a small subset of transcripts. Controversially, Limma associated better performance to more replicates. In the case of DS workflows, we found that DEXSeq achieved the highest TPRs percentages, followed by LimmaDS. Both of those found genes under DS and DS combined with changes in isoform expression levels, with better results in the latter. Those pipelines also showed higher percentages when more replicates were used. In both, DS and DIEDS groups, the best results were found for genes with the largest change in the major isoform expression. Finally, we suggest that if the number of replicates per condition is low, the workflows based on the Limma R package could be used to detect DIE and DS with high precision. The use of DESeq2 and DEXSeq workflows might be preferred when a high number of genes/isoforms are expected. If the number of replicates per condition is higher, we recommend the use of NOISeq workflow, for DIE analysis combined with any of LimmaDS or DEXSeq pipelines for DS analysis.

## KEY POINTS

- A number of workflows have been developed to either analyse differential gene or transcript expression and differential splicing using RNA-seq data. However, there is no clear consensus about the best practices for the simultaneous exploration of both types of transcriptional regulation. Our work analysed nine different pipelines or workflows to provide guidelines.
- The workflows choice directly impacts on the number of detected differential features (isoforms or genes) and the sensitivity and precision of the result.
- The number of isoforms per gene and the magnitude of the expression change influence the power of true detections. Fewer isoforms per gene and larger expression changes favour the detection of true positive differential features.
- The number of replicates and the amount of expected differentially expressed genes/isoforms between conditions should be taken into account when selecting the analysis pipeline.
- Workflows based on DESeq and DEXSeq are recommended for experiments with few heterogeneous samples; whereas, Limma and NOISeq pipelines will return better overall results when more replicates are available.

## AUTHORS DESCRIPTION

**Gabriela A. Merino** is a PhD student in Engineering Sciences and a Professor of Mathematical Analysis II at the National University of Cordoba, Argentina. Her doctoral research focuses on bioinformatics and statistical analysis of the next generation of sequencing data and is funded by the National Council of Science and Technology of Argentina (CONICET).

**Ana Conesa** is a bioinformatician and computational biologist developer of many different bioinformatics tools for functional annotation and analysis of transcriptomics data, quality assessment of short/long read sequencing data and multiomics integration. She enjoys a shared position as Head of the Genomics of Gene Expression Lab at the Centro de Investigación Príncipe Felipe (Valencia, Spain), and as Professor Bioinformatics at the Microbiology and Cell Sciences Department of the University of Florida at Gainesville, FL USA.

**Elmer A. Fernández.** Independent Researcher at National Council Of Science and Technology of Argentina (CONICET). He is a Biomedical Engineer from the National University of Entre Rios and he got his PhD in Advanced Computing and Artificial Intelligence from the University of Santiago de Compostela, Spain, He currently leads the Bioscience Data Mining Group at CIDIE-CONICET-UCC in Córdoba Argentina. He also leads de BioDataMining node of the National Bioinformatic Platform.

